# Bioinformatic characterisation of the effector repertoire of the strawberry pathogen *Phytophthora cactorum*

**DOI:** 10.1101/321141

**Authors:** Andrew D. Armitage, Eric Lysøe, Charlotte F. Nellist, Laura A. Lewis, Liliana M. Cano, Richard J. Harrison, May Bente Brurberg

**Author notes:** Address correspondence to Richard Harrison,.

## Abstract

The oomycete pathogen *Phytophthora cactorum* causes crown rot, a major disease of cultivated strawberry. We report the draft genome of *P. cactorum* isolate 10300, cultured from symptomatic *Fragaria* x *ananassa* tissue. Our analysis revealed that there are a large number of genes encoding putative secreted effectors in the genome, including nearly 200 RxLR domain containing effectors, 77 Crinklers (CRN) grouped into 38 families and numerous apoplastic effectors, such as phytotoxins (PcF proteins) and necrosis inducing proteins. As in other *Phytophthora* species, the genomic environment of many RxLR and CRN genes differed from core eukaryotic genes, a hallmark of the two-speed genome. We found genes homologous to known *Phytophthora infestans* avirulence genes including *Avr1, Avr3b, Avr4, Avrblb1* and *AvrSmira2* indicating effector sequence conservation between *Phytophthora* species of Clade 1A and 1C. The reported *P. cactorum* genome sequence and associated annotations represent a comprehensive resource for avirulence gene discovery in other *Phytophthora* species from Clade 1 and will facilitate effector informed breeding strategies in other crops.

## Introduction

The oomycetes are a diverse class of eukaryotic microorganisms that include pathogens of plants, animals and fungi (Simão et al. 2015). The causal agents of plant diseases are well represented in this phylogenetic class, with over 60% of known oomycetes characterised as plant pathogens (Thines and Kamoun 2010). Of these, the *Phytophthora* genus is responsible for some of the most economically and culturally significant diseases, including potato late blight caused by the pathogen *Phytophthora infestans*, stem rot of soybean caused by *Phytophthora sojae*, Sudden Oak Death caused by *Phytophthora ramorum* and blight of peppers and cucurbits caused by *Phytophthora capsici* (Kamoun et al. 2015).

The hemi-biotrophic oomycete pathogen *Phytophthora cactorum* (Lebert and Cohn) was identified as the causal agent of strawberry crown rot disease in 1952 (Eikemo et al. 2010) and is now considered a major disease of strawberry in temperate regions, leading to plant losses of up to 40% (Stensvand et al. 1999). *P. cactorum* is homothallic and produces oospores (resting spores) in diseased plant tissue. These can persist in the soil for many years and are an important source of inoculum in field production systems. *P. cactorum* is also a problem in the propagation of plants, risking rapid spread of the disease upon distribution (Fennimore et al. 2008). Chemical control via soil fumigation with chloropicrin 1,3-dichloropropene, dazomet and methyl bromide have proved effective in management of the pathogen (Cal et al. 2004). However, the phasing out of chemical fumigants in accordance with stricter European regulations (e.g. 91/414/EEC), has led to increased incidence of historically well-controlled soil-borne diseases. This has elevated the importance of integrating disease resistance into modern breeding germplasm. However, the functionality and durability of resistance is determined by pathogen encoded secreted effector proteins that can alter plant processes to aid infection (Franceschetti et al. 2017). Genome sequencing of *Phytophthora* spp. pathogens and subsequent functional characterisation of putative effector candidates from predicted gene models has provided a framework for study of *Phytophthora* diseases (Tyler et al. 2006; Haas et al. 2009; Lamour et al. 2012).

For *P. infestans*, characterization of effector genes, including the study of their interaction with host resistance genes (Armstrong et al. 2005; van Poppel et al. 2008; Oh et al. 2009; Champouret et al. 2009; Gilroy et al. 2011; Fry et al. 2015) has provided information about the durability of specific potato resistance genes. Similar suites of resistance genes have been identified against the soybean pathogen *P. sojae*, with fourteen major resistance genes at eight genomic loci determining a race structure within *P. sojae* (Sugimoto et al. 2012). This highlights the importance of understanding pathogen populations in the field and the associated genetic variation in effector complements. In contrast to these *Phytophthora* pathosystems, strawberry resistance to *P. cactorum* appears to be quantitative (partial; Denoyes-Rothan et al. 2004; Shaw et al. 2006; Shaw et al. 2008; Nellist et al. 2018), and no race structure has been reported to date. As such, resistance is not determined by a single gene-for-gene recognition, as often associated with RxLR (arginine, any amino-acid, leucine, arginine) effectors (Anderson et al. 2015). In soybean, quantitative resistance is observed alongside race-specific resistance and has been linked to the accumulation of PR1a (a matrix metalloproteinase), a basic peroxidase and a β-1,3-endoglucanase at the inoculation site (Vega-Sánchez et al. 2005) and to the accumulation of suberin in the roots (Thomas et al. 2007). For this reason, a range of effector candidates need to be considered when studying quantitative resistance in the strawberry pathosystem.

*Phytophthora* produce apoplastic effectors that are secreted to the extracellular space of the host and cytoplasmic effectors that are translocated to the host cytoplasm or intracellular compartments. Cytoplasmic RxLR’s are typified by an N-terminal signal peptide sequence allowing secretion of the protein, followed by an RxLR-EER motif that may be cleaved prior to secretion (Wawra et al. 2017), and a variable C-terminal domain, often containing WY domains (Win et al. 2012). RxLR effectors typically modulate host defense by suppressing host cell death (Anderson et al. 2015). The recognition of the RxLR class of effectors is mediated by a range of plant resistance proteins (Armstrong et al. 2005; Oh et al. 2009; Kourelis and van der Hoorn 2018).

Another major class of cytoplasmic effectors in *Phytophthora* pathogens are the Crinklers (CRN, for CRinkling and Necrosis), named due to the response observed when *P. infestans* CRNs were ectopically expressed in plants (Torto et al. 2003). CRNs have been shown to promote Pattern-Triggered Immunity (PTI), a process that is suppressed by RxLR effectors, indicating their functions may be associated with the necrotrophic stage of a hemi-biotrophic lifecycle (Stam et al. 2013b; Jupe et al. 2013; Stam et al. 2013a). Resistance is yet to be shown to this family of effectors but evidence has been presented for a heightened resistance response in *Nicotiana benthamiana* when infected with a *P. sojae* mutant overexpressing *PsCRN161* and in tomato plants infected with PVX vector containing *P. infestans crn2* (Torto et al. 2003; Rajput et al. 2015). CRNs characteristically possess an N-terminal LxLFLAK-motif connected with translocation, a DWL domain and the conserved recombination HVLVVVP-motif C-terminal domain. In some cases a DI domain is present between the LFLAK and DWL domain (Haas et al. 2009; Stam et al. 2013b). Functional studies have shown the LFLAK domain to be involved in entry into the host cell, following this CRNs target host nuclear processes, but the mechanisms of trafficking into the nucleus remain unknown (Amaro et al. 2017). Interestingly, most CRN effectors do not have predicted signal peptides or if they do, they exhibit lower SignalP scores (HMM models; Hidden Markov Model) compared to the RXLRs proteins. These weak *in silico* predictions of signal peptides in CRN proteins could be due to the CRNs being non-functional or due to non-classical methods of secretion from the pathogen (Amaro et al. 2017).

A diverse range of other secreted effectors are deployed during infection by *Phytophthora* spp. including plant cell wall degrading enzymes, protease inhibitors and phytotoxins of the PcF Toxin Family (Tian et al. 2005; Orsomando et al. 2011; Blackman et al. 2014). Furthermore, secreted non-effector proteins have been implicated in triggering HR in non-host species, such as elicitin INF1 (Kamoun et al. 1997). Elicitins are secreted sterol binding and carrier proteins, an essential protein family for *Phytophthora* spp., which are unable to produce sterols themselves due to an inability to produce oxidosqualene (Gottlieb et al. 1978; Wood and Gottlieb 1978).

With this work we aim to develop new genetic resources tools for the study of *Phytophthora* crown rot disease on cultivated strawberry including the first strawberry pathogen genome for *P. cactorum*, as well as the identification of candidate effectors from apoplastic and cytoplasmic families. *P. cactorum* has a diverse host range, infecting over 200 plant species (Erwin and Ribeiro 1996). This includes beech, for which a draft genome assembly was recently released (Grenville-Briggs et al. 2017). New approaches to identify *Phytophthora* CRNs are described, including their use to identify novel CRN families in *P. cactorum*, as well as highlighting additional CRNs in reference *Phytophthora* spp. genomes. These data represent valuable new resources for studying host adaptation within *P. cactorum* and enable the study of effector complements within *P. cactorum* and their comparison to Clade 1 *Phytophthora* spp. *P. infestans* and *P. parasitica*, as well as the more distant species *P. sojae* and *P. capsici* (Martin et al. 2014).

## Materials and Methods

### Pathogen isolate

*P. cactorum* Bioforsk isolate 10300 was sourced from symptomatic *Fragaria* x *ananassa* tissue, in Ås, Norway in 2006. Routine culturing was performed on V8 media at 20 °C. Genomic DNA was extracted from mycelium cultured in liquid Plich medium, using the OmniPrep™ kit for high quality genomic DNA extraction, following the manufacturer’s protocol.

### Pathogen sequencing and genome assembly

Genomic libraries from *P. cactorum* isolate 10300 were prepared for Illumina short read sequencing with insert sizes of 300 bp, 1 kb and 5 kb. Libraries with inserts of 300 bp and 1 kb were prepared using Illumina Truseq LT (FC-121–2001), whereas 5 kb mate-pair genomic libraries were prepared using Nextera Mate Pair gel-plus and gel-free protocols. Illumina sequencing was performed on the libraries using 2 × 75 bp reads for 300 bp and 1 kb insert libraries and 2 × 300 bp reads for the 5 kb insert library. Sequencing resulted in 42.86, 57.76 and 10.84 million reads from the 300 bp, 1 kb and 5 kb insert libraries, respectively. Removal of low quality and adapter sequences using fastqmcf, resulted in 41.29, 51.15 and 4.17 million reads from the 300 bp, 1 kb and 5 kb insert libraries, respectively.

*De-novo* genome assembly was performed using ABySS software version 1.3.7 (Simpson et al. 2009), using a kmer length of 53 bp. Contigs shorter than 500 bp were discarded and assembly statistics of remaining contigs were summarised using QUAST software version 3.0 (Gurevich et al. 2013). BUSCO software version 3.0.2 was used to assess the completeness of the assembly using the associated dataset of 303 core eukaryotic genes (CEGs) as a database for BUSCO analyses (Simão et al. 2015). RepeatModeler software version 1.0.8 and RepeatMasker software version 4.04 were used to identify repetitive elements and low complexity regions within the genome assembly (available at: http://repeatmasker.org).

### Gene models and ORF prediction

Gene prediction was performed on the softmasked *P. cactorum* genome using BRAKER1 software version 2 (Hoff et al. 2016), a pipeline for automated training and gene prediction of AUGUSTUS version 3 (Stanke and Morgenstern 2005). Evidence for gene models was generated using publicly available *P. cactorum* RNAseq reads (Chen et al. 2014), which were downloaded and aligned to *P*. *cactorum* assembly using STAR software version 2.5.3a (Dobin et al. 2013). Gene models were also called using CodingQuarry software version 2.0 (Testa et al. 2015), which was run using the “pathogen” flag parameter. CodingQuarry gene models were used to supplement BRAKER1 gene models, when individual CodingQuarry gene models were predicted in intergenic regions between BRAKER1 gene models.

Gene models were also supplemented with additional effector candidates from open reading frames (ORFs) located in intergenic regions of BRAKER1 and CodingQuarry genes. In addition, ORFs were predicted by translating sequences following all start codons in the genome until a stop codon or the end of the contig was reached. ORFs were predicted from sequences without any N’s and were translated to between 50–250 amino acids (aa) in length. All ORFs encoding proteins were screened for secretion signals followed by RxLR and CRN effector motifs (as described below) and of those testing positive, the ones present in intergenic regions were incorporated into gene models.

### Functional annotation of gene models

Draft functional annotations were determined for gene models using InterProScan-5.18–57.0 (Jones et al. 2014) and through identifying homology between predicted proteins and those contained in the SwissProt database (Bairoch and Apweiler 2000) using BLASTP (E-value > 1 × 10^-100^) (Altschul et al 1990). Homology was identified between predicted gene coding sequence and the Pathogen-Host Interactions database (PHIbase; www.phi-base.org/) (Winnenburg et al. 2006) using BLASTX (E-value > 1 × 10^-30^). Homology was also identified against a set of 50 previously characterised oomycete effectors/avirulence genes using BLASTN (E-value > 1 × 10^-30^). Functional annotation also identified the Carbohydrate-Active enZyme (CAZyme) encoding genes of *P. cactorum*. This was done using dbCAN (Huang et al. 2018) and using the CAZyme database classification (Cantarel et al. 2009).

Genes encoding putative secreted proteins were identified through prediction of signal peptides using SignalP software versions 2.0, 3.0 and 4.1 (Käll et al. 2004). Use of SignalP v2.0, as well as limiting secreted proteins to those with HMM scores greater than 0.9 and with cleavage sites between the 10^th^ and 40^th^ amino acid were consistent with previous RxLR prediction methodologies (Bhattacharjee et al. 2006; Win et al. 2007). Transmembrane proteins and membrane anchored proteins were identified using TMHMM version 2.0 and the GPI-SOM web-server, respectively (Krogh et al. 2001; Fankhauser and Mäser 2005). Proteins were considered ‘putatively secreted’ if they tested positive for a secretion signal using SignalP and lacked a transmembrane domain or membrane anchor signal. Additionally, Phobius software version 1.01 was used to screen proteins for secretion signals missed by SignalP (Käll et al. 2007). Proteins containing transmembrane domains or GPI anchored proteins were not excluded from the CRN and RxLR effector annotation pipelines discussed below.

### Crinkler effector identification

HMM models for CRN prediction were trained from CRN effectors predicted for *P. infestans, P. sojae, P. ramorum* and *P. capsici* (Tyler et al. 2006; Haas et al. 2009; Stam et al. 2013b). A HMM model training set of 271 (of 315) *Phytophthora* spp. CRNs were selected based on proteins identified in *P. infestans* and 65 (of 84) and *P. sojae* CRNs (Haas et al. 2009; Stam et al. 2013b). 63 excluded proteins lacked characteristic LFLAK or HVLVVP motifs from the LFLAK or DWL domains or contained ambiguous sites (‘X’s) in their sequence. The remaining proteins were considered to represent high confidence CRNs. Alignment of these sequences allowed training of a model to the LFLAK domain (from the conserved ‘MV’ to ‘LFLAK’ motifs) and a second model to the DWL domain (from the conserved ‘WL’ to the ‘HVLVVVP’ motifs). Putative CRNs were identified in predicted proteomes and translated ORFs by HMM searches using both LFLAK and DWL HMM models. Sequences required a HMM score greater than 0 for both models to be considered a putative CRN.

All predicted ORFs from the *P. cactorum* genome were screened using the trained LFLAK and DWL HMM models. Those ORFs with an HMM score greater than 0 for both FLAK and DWL HMM models were retained. As some of these ORFs were found to overlap, redundancy was removed from the dataset by retaining only the ORFs with the greatest LFLAK domain HMM score. The putative CRN ORFs located in intergenic regions of BRAKER1/CodingQuarry gene models were integrated into the final set of gene models.

### RxLR effector identification

Motif and HMM based approaches were used to predict genes encoding RxLR proteins in *P. cactorum* and reference *Phytophthora* spp. genomes. Motif based prediction was built upon previous N-terminal RxLR identification pipelines (Torto et al. 2003). Secreted proteins were considered putative RxLRs if an RxLR motif was present up to 100 aa downstream of the signal peptide cleavage point and the protein carried an EER motif within 40 aa downstream of the RxLR position. EER motifs were searched for using the Python regular expression ([ED][ED] + [KR]).

Heuristic based methods for RxLR prediction were used to complement motif presence methods. A previously described RxLR HMM model was used to statistically assess secreted proteins for the presence of N-terminal RxLR-like regions (Whisson et al. 2007). Hits with an HMM score greater than 0 were considered putative RxLR proteins.

All predicted ORFs carrying a secretion signal in the *P. cactorum* genome were screened for RxLR motifs and homology to HMM models. As some predicted ORFs were found to overlap one another, redundancy was removed from the dataset retaining only the ORF with the greatest SignalP HMM score. Those RxLR-containing ORFs located in intergenic regions of

BRAKER1/CodingQuarry gene models were integrated into the final set of gene models. All RxLR candidates were screened for presence of C-terminal WY-domains using a previously described HMM model (Win et al. 2012).

### Gene orthology analysis

Ortholog identification was performed using OrthoFinder software version 1.1.10 (Emms and Kelly 2015) on all *P. cactorum* isolate 10300 predicted proteins and the proteomes of publicly available *Phytophthora* species *P. infestans, P. parasitica, P. capsici* and *P. sojae*. Venn diagrams were plotted using the R package software version 3.5.2, VennDiagram package version 1.6.20 (Chen and Boutros 2011). Further clustering was performed on the combined set of CRN effector proteins from *P. cactorum, P. infestans, P. parasitica, P. capsici* and *P. sojae* using OrthoMCL software version 2.0.9 (Li et al. 2003), with the inflation value set to 5 in order to increase resolution within groups.

## Results

### Pathogen genome assembly

*De-novo* genome assembly of the Norwegian strawberry *P. cactorum* isolate 10300 using ABySS (Simpson et al. 2009) generated a 59.3 Mb assembly in 4,623 contigs, with an N50 value of 56.3 kb (Table 1). Total assembly size was smaller than the other available *Phytophthora* spp. assemblies including *Phytophthora* Clade 1C relatives *P. parasitica* and *P. infestans* assemblies, but was found to contain a similar or greater gene space within the assembly, with BUSCO identifying 283 of 303 CEGs. Of these CEGs, 274 were present in a single copy within the assembly. The *P. cactorum* genome was found to be repeat-rich, with RepeatModeler and RepeatMasker identifying 18% of the genome as repetitive or low complexity regions. This level of repetitive content was considerably lower than observed in *P. infestans*, but comparable to *P. capsici* that has a similarly sized genome of 64 Mb. Meaningful comparisons of repeat content could not be made between the *P. cactorum* and *P. parasitica* genomes as the scaffolded *P. parasitica* assembly contained a high percentage of N’s (Table 1). A total of 23,884 genes encoding 24,189 proteins were predicted from the *P. cactorum* genome with 21,410 genes predicted from the BRAKER1 pipeline (Hoff et al. 2016), 2,434 additional genes from CodingQuarry (Testa et al. 2015), and a further 40 coding genes from intergenic ORFs identified as putative secreted RxLR or CRN effectors. The number of predicted genes reported in *Phytophthora* spp. shows considerable variation between studies, however *P. cactorum* gene models contained the greatest number of complete single copy CEGs among the assessed *Phytophthora* spp., indicating good representation of gene space within gene models (Table 1).

**Table 1:** Assembly and gene prediction statistics for the *Phytophthora cactorum* genome, with reference to publicly available *Phytophthora* spp. genomes from Clades 1, 2 and 7 (Blair et al. 2008). Number of core eukaryotic genes (CEGs) identified as complete and present in a single copy are shown for each genome/set of gene models, as determined by BUSCO.

| Species | *P. cactorum* | *P. parasitica* | *P. infestans* | *P. capsici* | *P. sojae* |
| --- | --- | --- | --- | --- | --- |
| Phylogenetic clade | 1a | 1b | 1c | 2 | 7 |
| Strain | 10300 | INRA-310 | T30-4 | LT1534 | P6497 |
| Assembly size (mb) | 59.3 | 82.4 | 228.5 | 64 | 83 |
| Number of contigs | 4623 | 708 | 4921 | 917 | 83 |
| Number of contigs (>1 kb) | 2913 | 708 | 4598 | 917 | 83 |
| Largest contig (kb) | 301 | 4,724 | 6,928 | 2,170 | 13,391 |
| N50 (kb) | 56.3 | 888 | 1,589 | 706 | 7,609 |
| N's per 100 kb | 4006 | 34,613 | 16,806 | 12,466 | 3959 |
| Repeatmasked (mb) | 10.8 (18%) | 7.0 (8%) | 152.1 (67%) | 13.6 (21%) | 23.7 (29%) |
| CEGs in the assembly | 274 (90%) | 271 (89%) | 255 (84%) | 269 (89%) | 270 (89%) |
| Predicted genes | 23,884 | 20,822 | 17,787 | 19,805 | 26,584 |
| CEGs in gene models | 272 (89%) | 271 (89%) | 257 (85%) | 261 (89%) | 262 (86%) |

### Orthology analysis

Clustering of predicted proteins from the five *Phytophthora* spp. using OrthoFinder, resulted in 15,162 orthogroups containing 95,739 proteins (87.7% of the total). A total of 20,157 (84%) of predicted *P. cactorum* proteins had identified orthologs in other *Phytophthora* spp. Of these, 9,553 orthogroups contained proteins from all five species, with 6,767 orthogroups consisting of a single protein from each species (Fig. 1).

**Figure 1:**
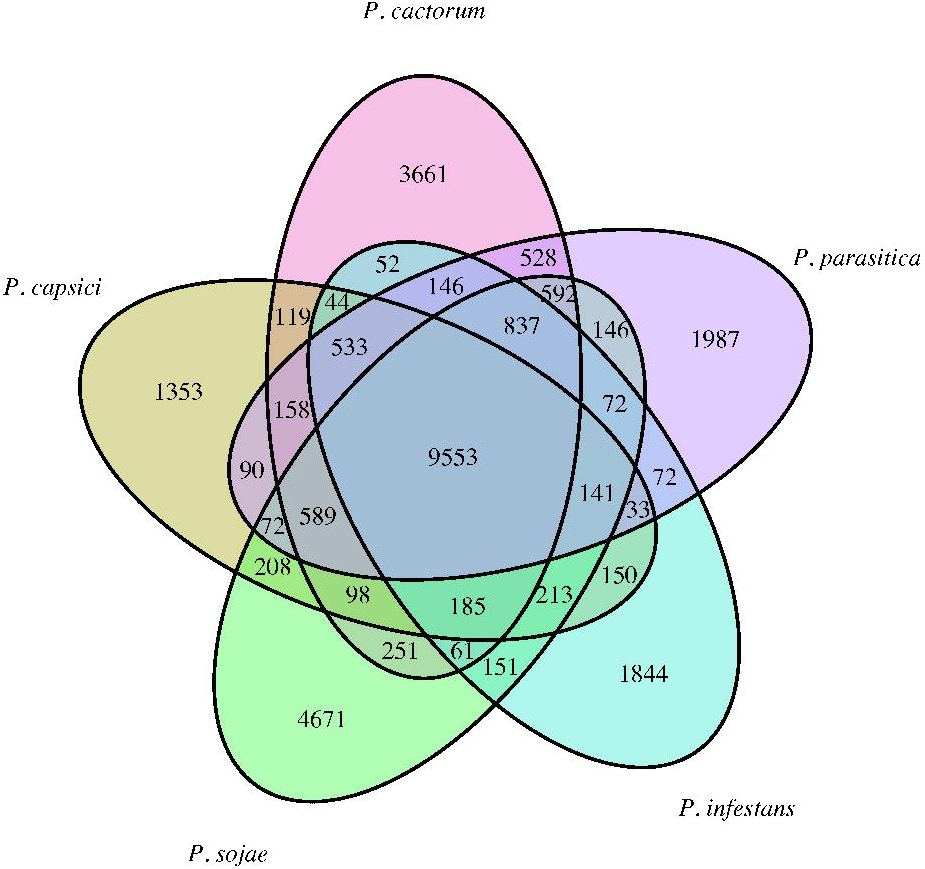
Number of shared and unique ortholog groups between *Phytophthora* spp. Orthogroups determined from clustering 109,187 proteins from *P. cactorum, P. parasitica, P. infestans, P. capsici* and *P. sojae*.

### Intergenic distance

Intergenic distance (IGD) was determined for each gene, by counting the number of bp to the nearest gene in 5’ and 3’ directions. Genes that were on the end of a contig and therefore did not have a neighbouring gene up- or down-stream were discarded from this analysis.

### Functional annotation and secretome prediction

Genomic locations of *P. cactorum* 10300 gene models, their orthology assignment and predicted functional annotations are summarised in Supplementary Data 1. A total of 2,234 genes encoded putatively secreted proteins. The number of predicted genes encoding secreted MAMPs, apoplastic effectors and cytoplasmic effectors are summarised in Table 2 and discussed below.

**Table 2:** Total number of predicted effector gene candidates in *Phytophthora cactorum* isolate 10300 and genes associated with triggering plant basal defence (microbe associated molecular patterns, MAMPs). Numbers of genes shown relate to genes encoding predicted secreted proteins.

| Category | Family | Number of proteins |
| --- | --- | --- |
| MAMP | Sterol binding proteins | 47 |
|  | Transglutaminase proteins | 15 |
| Apoplastic effectors | Secreted CAZymes | 282 |
|  | Protease inhibitors (glucanase) | 2 |
|  | Phytotoxins | 2 |
|  | Necrosis inducing proteins | 24 |
|  | Cutinases | 4 |
|  | Protease inhibitors (kazal) | 14 |
|  | Protease inhibitors (cathepsin) | 3 |
|  | Protease inhibitors (cystatin) | 3 |
| Cytoplasmic effectors | Crinklers | 77 |
|  | RxLRs | 199 |

## Microbe associated molecular pattern (MAMP) genes

### Sterol-binding proteins

*Phytophthora* spp. lack the ability to synthesize sterols and are reliant on assimilation from the environment. Secreted sterol binding proteins are known Microbe-Associated Molecular Patterns (MAMPs), triggering host recognition. For this reason, they are also referred to as “elicitins”. A total of 66 genes possessed an elicitin domain (IPR002200), of which 47 were predicted as secreted. These genes showed high levels of local duplication, with 41 of the 66 genes in 11 elicitin gene clusters.

### Transglutaminase proteins

The *P. sojae* cell wall glycoprotein GP42 is an elicitor of host defence and is functionally characterized as a Ca^2+^ –dependant transglutaminase (Brunner et al. 2002). Recognition of the protein by plant hosts is lost upon mutation of the transglutaminase domain, indicating its importance for recognition. A total of 23 *P. cactorum* genes were predicted to encode transglutanimase domains (IPR032048). These were distributed through 10 orthogroups, with 13 proteins contained in a single orthogroup (OG0000097). BLAST searches identified 19 *P. cactorum* genes with homology to *P. sojae* GP42 (Supplementary Data 2), each of which was identified by domain searches. Of the 23 proteins carrying transglutaminase domains, 15 were predicted to be secreted.

## Apoplastic effectors

### Carbohydrate active enzymes (CAZymes)

CAZymes play a direct role in pathogenicity, contributing to plant cell wall degradation. A total of 696 transcripts encoding CAZymes were identified in the *P. cactorum* 10300 genome, of which 352 were predicted as carrying an N-terminal signal peptide and 282 were predicted as secreted (removing those with transmembrane and GPI anchors domains). These secreted CAZymes were distributed through glyceraldehyde hydrolases (GH), carbohydrate binding molecules (CBM), auxiliary activity (AA), carbohydrate esterase (CE) and pectin lyase (PL) and glycosyl transferases (GT) families containing 172, 22, 6, 24, 37 and 21 proteins, respectively.

The profile of *P. cactorum* cell wall degrading enzymes was investigated through further study of GH, CBM, AA, CE and PL families (Table 3). Substrate specificity was not further investigated within the GT proteins due to wide polyspecificity (multiple substrates associated with the same GT family) within this group. Cell wall degrading enzymes can be summarized by functions, targeting cellulose, hemicellulose or pectin. Cellulase activity is represented in seven GH families, two CBM families and three AA families. Cellulases are well represented in the CBM compliments of *P*. *parasitica, P. ramorum* and *P. sojae*, where CBM1 and CMB63 represented the two largest groups of CBMs. This was also true for *P. cactorum*, where CBM63 and CBM1 proteins represented 81% of the putatively secreted CBM molecules. This is in contrast to fungal necrotrophs which typically possess 1–3 CBM3 proteins (Blackman et al 2014). In fungi, CBM1 and CBM63 domains are predominantly accompanied by additional modules (Blackman et al 2014), however none of the CBM63 or CBM1 proteins in *P. parasitica* are accompanied by other catalytic modules (Blackman et al 2014). This was also true of *P. cactorum* CBM63 or CBM1 CAZymes. Hemicellulose targeting secreted CAZYmes were represented in 12 GH families, one CBM family and four CBM families. The *P. cactorum* genome encodes large numbers of proteins involved in pectin modification, including GH groups GH28 and GH81 representing the third and fifth most abundant GH groups (15 and 12 proteins), CE8 representing the most abundant CE group (8 proteins) and 37 proteins from PL families PL3, PL1 and PL4. *Phytophthora* spp. are reported to carry expanded pectin targeting CDWE in comparison to fungi (Blackman et al. 2014). In total, 79 putatively secreted CWDE targeted pectin, which is comparable to the 86 predicted in *P. parasitica*, and in contrast to fungi, which typically have less than 20 PL proteins (Blackman et al. 2014).

Secreted enzymes targeting β-1,3-glucan targeting CAZymes may function in breakdown of callose, as deposited by the host upon triggering of basal defense. β-1,3-glucans are also found in the pathogen, being present in the oomycete cell wall where they act as MAMPs triggering plant basal defence (Robinson and Bostock 2015). Reflecting this, *P. cactorum* carried a large number of genes (31) encoding putatively secreted proteins from five different families targeting β-1,3-glucan. Notably, 21 genes encoded GH17 proteins, which was the most abundant CAZyme family.

**Table 3:** Profile of secreted *Phytophthora cactorum* Carbohydrate-Active enZymes (CAZymes) from glyceraldehyde hydrolase (GH), carbohydrate binding molecules (CBM), auxiliary activity (AA), carbohydrate esterase (CE) and pectin lyase (PL) families, as identified by dbCAN. Numbers are shown for total numbers of N-terminal signal peptide containing proteins, and those considered putative secreted proteins, which lack transmembrane signals of membrane anchors. Target substrates of for each family is shown. HG = homogalacturonan, RGI = rhamnogalacturonan I; GlcNAc = N-acetylglucosamine.

| CAZY family | Substrate | Signal peptide | Secreted proteins |
| --- | --- | --- | --- |
| GH17 | β-1,3-glucans | 21 | 21 |
| GH3 | cellulose, hemicellulose (xyloglucans), pectin (RGI), AGPs | 19 | 16 |
| GH28 | pectin (HG) | 15 | 15 |
| GH16 | hemicellulose (xyloglucans), β-1,3-glucans | 18 | 12 |
| GH81 | pectin (RGI) | 12 | 12 |
| GH30 | cellulose, hemicellulose (xyloglucans), pectin (RGI), AGPs | 12 | 11 |
| GH12 | cellulose, hemicellulose (xyloglucans) | 12 | 9 |
| GH72 | β-1,3-glucans | 10 | 8 |
| GH1 | cellulose, hemicellulose (xyloglucans), pectin (RGI) | 10 | 8 |
| GH5 | cellulose, hemicellulose (xyloglucans, galactomannans), β-1,3-glucans | 8 | 8 |
| GH6 | cellulose | 7 | 6 |
| GH78 | pectin (RGI) | 6 | 6 |
| GH43 | hemicellulose (xylans), pectin, AGP | 5 | 4 |
| GH31 | starch, hemicellulose (xyloglucans) | 5 | 4 |
| GH131 | β-1,3-glucans, hemicellulose (β-1,4-glucans) | 5 | 4 |
| GH7 | cellulose | 4 | 4 |
| GH53 | pectin (RGI) | 4 | 3 |
| GH32 | sucrose | 3 | 3 |
| GH19 | N-linked oligosaccharides | 3 | 3 |
| GH10 | hemicellulose (xylans) | 3 | 3 |
| GH17, CBM13 | β-1,3-glucans | 3 | 2 |
| GH18 | N-linked oligosaccharides | 2 | 2 |
| GH54 | pectin (RGI) | 1 | 1 |
| GH47 | N-linked oligosaccharides | 1 | 1 |
| GH38 | N-linked oligosaccharides | 1 | 1 |
| GH2 | hemicellulose (mannans), glycoproteins (mannans) | 1 | 1 |
| GH16, GT48 | hemicellulose (xyloglucans), β-1,3-glucans | 1 | 1 |
| GH13 | starch | 1 | 1 |
| GH105 | pectin (RGI) | 1 | 1 |
| GH31, CBM25 | starch | 1 | 1 |
| GH89 | N-linked oligosaccharides | 2 | 0 |
| GH114 | α-1,4-polygalactosamine | 1 | 0 |
| GH5, CBM43 | β-1,3-glucans | 1 | 0 |
| CBM63 | cellulose | 11 | 9 |
| CBM1 | cellulose | 10 | 9 |
| CBM47 | fucose binding | 2 | 1 |
| CBM9 | hemicellulose (xylans) | 1 | 1 |
| CBM36 | xylanase | 1 | 1 |
| CBM32 | galactose, PGA and β-galactosyl-β-1,4-GlcNAc | 1 | 1 |
| CBM38 | inulin binding | 1 | 0 |
| AA2 | lignin | 4 | 3 |
| AA8 | cellulose | 3 | 1 |
| AA10 | cellulose | 2 | 1 |
| AA9 | cellulose | 1 | 1 |
| AA7 | Glycolate oxidase | 2 | 0 |
| CE8 | pectin (HG) | 9 | 8 |
| CE1 | hemicellulose | 8 | 4 |
| CE10 | non-carbohydrate substrates | 6 | 3 |
| CE13 | pectin (HG) | 5 | 3 |
| CE5 | hemicellulose | 3 | 3 |
| CE12 | pectin (HG, RGI) | 2 | 2 |
| CE3 | hemicellulose | 1 | 1 |
| CE4 | hemicellulose, N-linked oligosaccharides | 1 | 0 |
| PL3 | pectin (HG, RGI) | 21 | 17 |
| PL1 | pectin (HG) | 19 | 16 |
| PL4 | pectin (RGI) | 4 | 4 |

### Glucanase inhibitors

Non-CAZyme proteins are involved in preventing host recognition of *Phytophthora* β-1,3-glucans. Glucanase inhibitor proteins (GIPs) are serine proteases that inhibit degradation of β-1,3/1,6- glucans in the pathogen cell wall and/or the release of defence-eliciting molecules by host endoglucanases (Kamoun 2006). These serine proteases contain a domain that shows homology to the chymotrypsin class of serine proteases, however they lack proteolytic activity and as such belong to a broader class of proteins called serine protease homologs (Damasceno et al. 2008). A total of 34 *P. cactorum* genes were predicted to encode proteins with chymotrypsin domains (IPR001314), with 24 of these predicted as secreted and 28 as homologs of GIP proteins from *P. infestans* and *P. sojae*. Three of the *P. cactorum* proteins were members of a single orthogroup containing *P. infestans* GIP proteins (PITG_13636, PITG_21456), of which two were predicted as secreted and therefore represent high-confidence glucanase inhibitor candidates.

### Phytotoxins

The PcF toxin family was first described from *P. cactorum* (Orsomando et al. 2001). BLAST searches identified g2968.t1 as homologous to PcF (NCBI accession: AF354650.1). This gene was a member of an orthogroup with two members from *P. infestans*, one member from *P. parasitica* and two members from *P. capsici*. InterProScan annotation identified two additional phytotoxin candidates (g10773.t1, g16798.t1) carrying the PcF domain (Pfam: PF09461) in addition to g2968.t1. Each of the three identified genes encoded a protein with a N-terminal secretion signal but g16782.t1 was also predicted to encode a transmembrane domain.

### Necrosis inducing proteins

Necrosis inducing proteins (NLPs) are produced by bacterial, fungal and oomycete plant pathogens (Gijzen and Nürnberger 2006). These proteins are associated with the transition from biotrophy to necrotrophy in *Phytophthora* spp. and act by triggering cell death (Feng et al. 2014). These proteins may also stimulate immune responses in the host. Forty three proteins carrying NLP-like domains (PF05630, IPR008701) were identified in *P. cactorum*, of which 24 were predicted as secreted. Thirty of these proteins were NPP1 homologs in PHIbase, of which 21 were predicted as secreted.

Twenty five of the 43 genes also showed homology to assembled *P. cactorum* transcripts from previous work (Chen et al. 2014). The 43 proteins were distributed through 16 orthogroups, including all 13 members of orthogroup 75 and all 12 members of orthogroup 12. Alignment of all proteins in the 16 NLP orthogroups showed that these proteins represent Type 1 NLPs, through conservation of two cytosine sites (alignment positions 624 and 661 in Supplementary Data 3).

### Cutinases

In addition to the plant cell wall, cutin acts as a barrier to host penetration by plant pathogens. Pathogens often employ methods to circumvent this barrier such as colonisation via stomata or through wounds. In total, seven genes were annotated as cutinase genes (PF01083), and four of these putative cutinases were predicted as secreted. Interestingly, three of the four secreted cutinases and a non-secreted cutinase (g10526, g10527, g10528, g10530) were clustered in a 5 kb region of the genome. Two of these genes belonged to the same orthogroup, which showed an expansion of genes in *P. sojae* (14 genes), but similar numbers in the other *Phytophthora* spp. (3–4 genes). The other two *P. cactorum* genes were present in single-gene orthogroups unique to *P. cactorum*. Closer investigation revealed that one of these two genes was truncated, and the other incomplete due to being located on the end of the contig.

### Protease inhibitors

Plant hosts secrete proteases into the apoplastic space to degrade pathogen-secreted effectors. As such, oomycetes are known to secrete protease inhibitors to counteract these defenses (Tian et al. 2005). A total of 22 genes encoding Kazal-type protease inhibitors (IPR002350) were identified in *P. cactorum* gene models, with 14 of these predicted as secreted. It was noted that 12 of the 22 genes were located within 8 kb of another Kazal-domain encoding gene, in clusters of two or three genes. Despite this, the 22 genes represented 18 different orthogroups, indicating historical duplication and divergence between these effector genes. A further four genes encoding proteins with cathepsin propeptide inhibitor domains (IPR013201) were identified, three of which were predicted as secreted. All were located on different contigs and were members of distinct orthogroups. A number of secreted cystatin-like cysteine protease inhibitors have been identified from *P. infestans* (EPIC1-EPIC4), including EPIC2B which has been shown to inhibit the tomato defence response through interaction with an apoplastic papain-like cysteine protease (Tian et al. 2007). Three *P. cactorum* genes were predicted to encode secreted cystatin-like cysteine protease inhibitors, containing cystatin (IPR000010, IPR027214) or cystatin protease inhibitor (IPR018073, IPR020381) domains. These genes were in three orthogroups, each containing a single gene from *P. cactorum*. BLAST searches identified the three genes as homologs of *EPIC1, EPIC3* and *EPIC4*.

## Cytoplasmic effectors

### Crinkler annotation

A novel method of CRN prediction was developed based upon identification of the characteristic LFLAK and DWL domains. Trained HMM models are provided in Supplementary Data 4. Application of the LFLAK DWL models to *P. infestans* and *P. capsici* was used to validate the LFLAK-DWL approach. In total, 265 *P. infestans* and 175 *P. capsici* proteins were predicted to encode putative CRNs. Of the 194 proteins previously identified as CRNs in *P. infestans* (Haas et al. 2009), 35 were not identified by the LFLAK-DWL approach, each lacking the ‘HVLVVVP’ motif from the DWL domain. Similar results were observed for results from *P. capsici*, where 71 of the 84 previously identified CRNs were identified by the LFLAK-DWL approach (Stam et al. 2013b), and the remaining 13 were found to contain ambiguous sites (‘X’s). Application of the LFLAK-DWL to reference gene models and ORFs allowed identification of 265 CRNs in *P. infestans*, 35 in *P. parasitica*, 114 in *P. capsici* and 159 in *P. sojae*, with 4, 98, 32 and 89 candidates identified from translated ORFs, respectively (Fasta sequences available in Supplementary Data 5).

Application of the developed LFLAK-DWL approach to *P. cactorum* identified a total of 77 putative CRN effector genes, with three of these identified from ORF gene models. Inspection of the *P. cactorum* CRN gene models showed that 17 (22%) were incomplete, lacking stop codons due to being located on the ends of contigs. This may reflect the modular structure and duplication of CRNs leading to difficulty in genome assembly of these regions. CRNs are known to be secreted from the host cell but often lack predictable secretion signals, e.g. 58% of identified *P. capsici* CRNs lacking secretion signals (Stam et al. 2013b). We found similar results with 56% of *P. cactorum* CRNs lacking a signal peptide as predicted by SignalP 2, 3, 4 and Phobius software. Phobius was more sensitive than SignalP 2, 3 and 4, identifying signal peptides in 32 of the 77 CRNs as secreted, whereas the SignalP approaches predicted a combined total of 22 as secreted, with two that were not detected by Phobius.

The modular structure of CRNs and the diversity of CRN domains within *Phytophthora* spp. was further investigated using an orthology analysis on the total set of 650 predicted CRNs between the five studied species. Clustering using orthoMCL resulted in 73 groups of CRN proteins, with groups observed to separate by C-terminal domain (Fig. 2). All of the 39 previously described C-terminal domains were identified within the clustered proteins, as well as the variable DI domain within the N-terminal region (Haas et al. 2009; Stam et al. 2013b). *P. cactorum* CRNs were present in groups representing 21 of these domains, whereas 14, 31, 31 and 27 domains were represented in groups containing *P. parasitica, P. infestans, P. capsici* and *P. sojae* CRNs. *P. infestans* showed signs of gene expansion in some groups including those encoding DXZ domains (59 *P. infestans* proteins vs 4–13 from other species), D2 domains (39 *P. infestans* proteins vs 1–8 from other species), DHB-DXX-DHA domains (23 *P. infestans* proteins vs 1–4 from other species) proteins. Similar expansion was not observed in *P. cactorum* CRN genes, with most populous groups representing DXZ, DN17 and DFA-DDB/DDC domains. Many proteins in *P. infestans* expanded orthogroups were identical to one another, indicating that CRN proteins are subject to frequent duplication, and as such the total numbers of CRNs observed in a genome is likely to be highly influenced by the quality of the genome assembly. An additional 137 predicted CRN proteins in 37 orthogroups did not contain any recognizable CRN domains.

**Figure 2:**
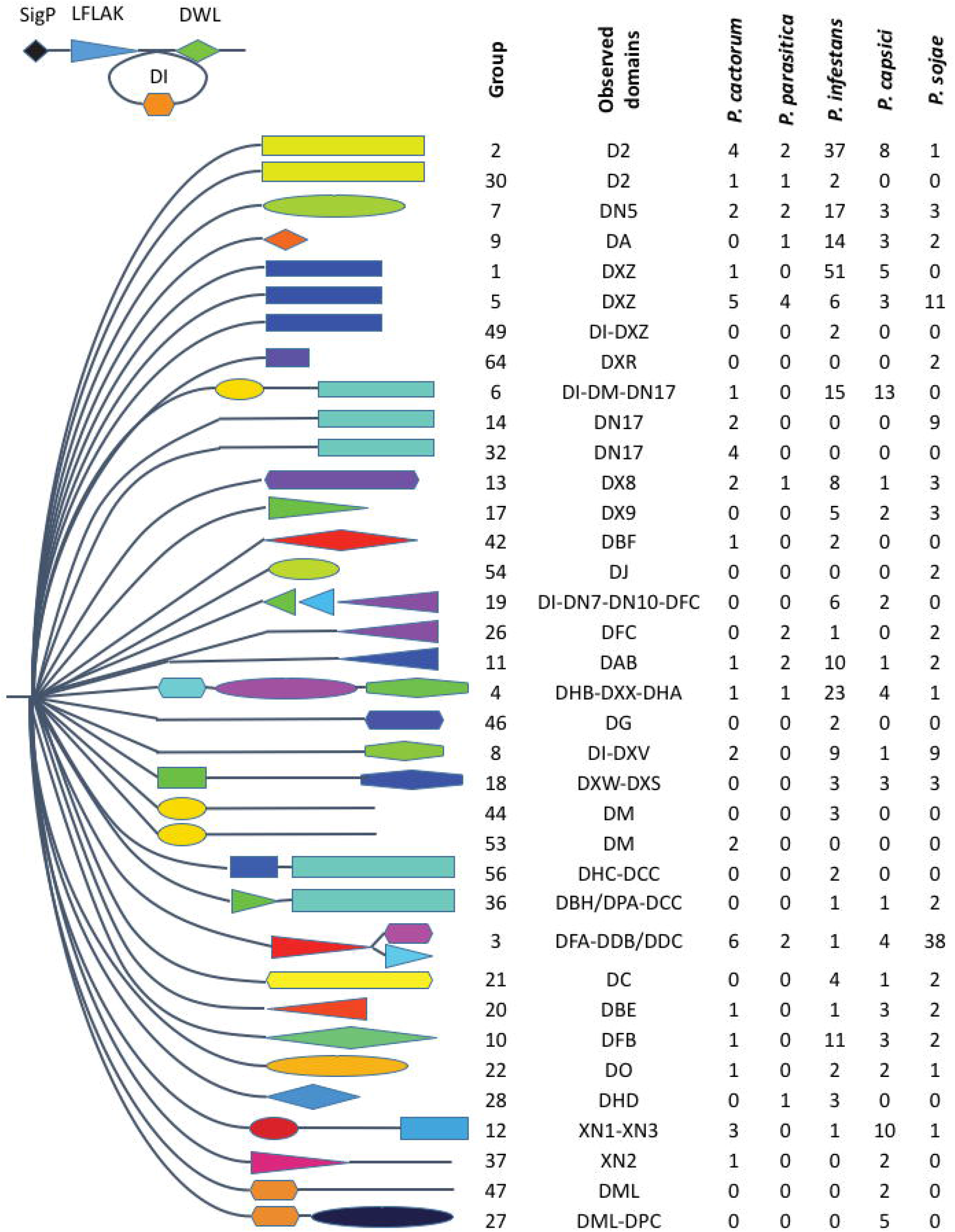
Clustering of *Phytophthora* spp. Crinklers (CRNs) separated by their C-terminal domain. All CRNs possessed a conserved LFLAK and DWL domain, some also possessed a DI domain in the N-terminal region. CRN proteins were observed to cluster by C-terminal domain as described in Haas et al. (2009) and Stam et al. (2013). The cluster (group) of proteins is shown along with observed domains and the number of *P. cactorum, P. parasitica, P. infestans, P. capsici* and *P. sojae* genes contained within each group.

### RxLR identification

A combined approach of regular expression searches for RxLR-EER motifs, as well as searches using HMM models identified 199 putative RxLR effectors in the *P. cactorum* assembly, with 162 of these predicted from predicted gene models and a further 37 from ORFs. Searches for WY domains found 92 WY-domain containing RxLRs. Functional annotation was largely absent for these proteins, but InterProScan annotations were present for ten proteins and a further five were predicted to be CAZymes (Table 4). Many of these domains have been associated with virulence in *Phytophthora* or other organisms (Sperisen et al. 2005; Xu et al. 2008; Bouwmeester et al. 2011; Blackman et al. 2014; Kong et al. 2015; Dong and Wang 2016). This included three RxLRs with Nudix-hydrolase annotations, a domain present in Avr3b. Avr3b from *P. sojae* is expressed at early stages of infection and delivered into the host cell where it maturates itself through recruitment of GGmCYP1, leading to suppression of effector triggered immunity (Dong et al. 2011; Kong et al. 2015). Genes in ortholog groups containing *PiAvr3b* and other characterised RxLRs were identified (Table 5). *P. cactorum* carried genes in orthogroups containing *P. infestans Avr1, Avr3b, Avr4, Avrblb1 and Avr-Smira2*. Avr1 is understood to manipulate basal defence through interaction with a plant exocyst subunit and thereby disturbing vesicle trafficking (Du et al. 2015). Two genes from *P. cactorum* were in the same orthogroup as *P. infestans Avr-blb1*, however one was truncated. Truncation has been observed in ∼10% of *P. sojae* and *P. ramorum* RxLRs (Jiang et al. 2008). Furthermore, truncation leading to loss of function in Avr4 has been shown to prevent host recognition, determining a race structure in *P. infestans* (van Poppel et al. 2008). Avr-blb1 is understood to interact with a lectin receptor kinase associated with the plasma membrane, leading to destabilising of the cell wall-plasma membrane to promote infection (Bouwmeester et al. 2011). A total of 35 *P. cactorum* RxLR candidates were members of orthogroups containing a single gene from both *P. cactorum* and *P. infestans*. Similar orthology assignments could be made for 33 *P. cactorum* RxLR candidates and *P. sojae* genes. Characterisation of these core RxLRs will aid understanding of the fundamental infection strategy conserved between *Phytophthora* spp.

**Table 4:**
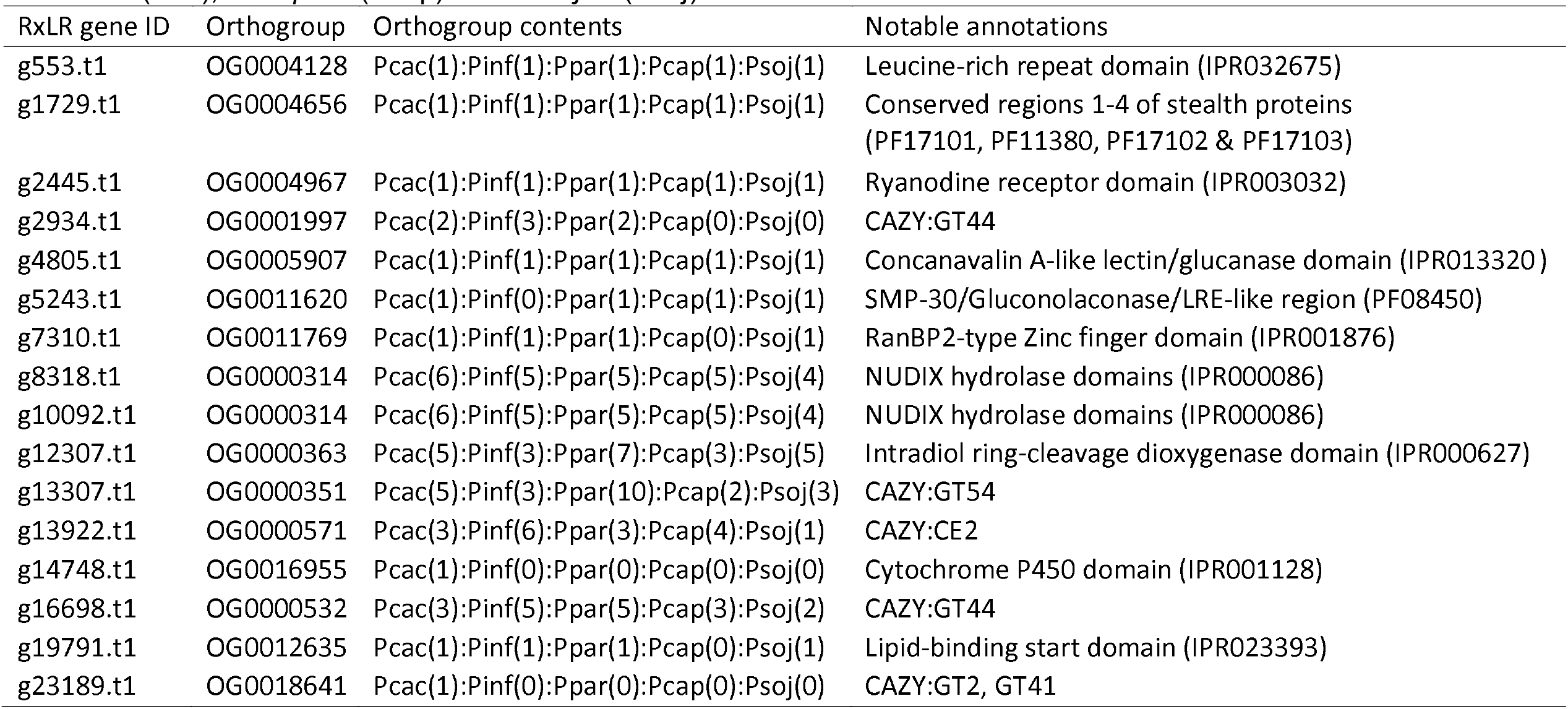
Functional annotations of *Phytophthora cactorum* RxLR candidates. Orthogroup assignment shows conservation of these genes throughout *Phytophthora* spp. Numbers of genes in each orthogroup are shown for *P. cactorum* (Pcac), *P. parasitica* (Ppar), *P. infestans* (Pinf), *P. capsici* (Pcap) and *P. sojae* (Psoj).

**Table 5:** *Phytophthora cactorum* genes in orthogroups shared with characterized *P. infestans* RxLR candidates. Orthogroup assignment shows conservation of these genes throughout *Phytophthora* spp. Numbers of genes in each orthogroup are shown for *P. cactorum* (Pcac), *P. parasitica* (Ppar), *P. infestans* (Pinf), *P. capsici* (Pcap) and *P. sojae* (Psoj).

| *P. cactorum*<br>gene ID | Contig | *P. infestans*<br>Avr gene | *P. infestans*<br>gene ID | Orthogroup | Orthogroup contents | Notes |
| --- | --- | --- | --- | --- | --- | --- |
| g15126.t1 | contig_485 | Avr1 | PITG_16663 | OG0000777 | Pcac(2):Pinf(2):Ppar(4):Pcap(4):Psoj(2) | TLLR at RxLR motif location |
| g16706.t1 | contig_608 | Avr1 | PITG_16663 | OG0000777 | Pcac(2):Pinf(2):Ppar(4):Pcap(4):Psoj(2) |  |
| g5545.t1 | contig_94 | Avr3b | PITG_15732 | OG0013112 | Pcac(1):Pinf(1):Ppar(1):Pcap(0):Psoj(0) | NUDIX hydrolase domain (IPR000086) |
| g4951.t1 | contig_80 | Avr4 | PITG_07387 | OG0011587 | Pcac(1):Pinf(1):Ppar(2):Pcap(0):Psoj(0) |  |
| g6635.t1 | contig_121 | Avrblb1 | PITG_21388 | OG0001713 | Pcac(2):Pinf(2):Ppar(4):Pcap(0):Psoj(0) | Truncated protein |
| g6663.t1 | contig_121 | Avrblb1 | PITG_21388 | OG0001713 | Pcac(2):Pinf(2):Ppar(4):Pcap(0):Psoj(0) |  |
| g15879.t1 | contig_543 | AvrSmira2 | PITG_07558 | OG0000427 | Pcac(2):Pinf(4):Ppar(5):Pcap(3):Psoj(7) |  |
| g18867.t1 | contig_844 | AvrSmira2 | PITG_07558 | OG0000427 | Pcac(2):Pinf(4):Ppar(5):Pcap(3):Psoj(7) |  |

Thirteen RxLR candidates lacked a recognisable EER motif and were not identified by the RxLR HMM model, but were identified by the presence of secretion signal, RxLR motif and WY domain. BLAST searches identified two of these genes as homologs to *P. infestans Avr-smira2* and a further four of these genes were identified as homologs to *P. sojae PSR2* and two as homologs to *Avh5*. Homologs to these characterised RxLR genes highlight the importance of using multiple sources of evidence in RxLR identification.

### Genomic distribution of *P. cactorum* effectors

Rapidly evolving RxLR and CRN genes are predominantly located in gene-sparse regions, with greater IGDs than core eukaryotic genes (Haas et al. 2009). The 5’ and 3’ flanking distance between each *P. cactorum* gene and its neighbours were taken as measurements of local gene density (Fig. 3), following exclusion of 5,041 genes (21%) that neighboured a contig break (Table 6). Effector genes were located in gene sparse regions of the *P. cactorum* genome, with RxLR genes having greater mean 5’ and 3’ IGDs than observed for non-RxLR genes (*p* < 0.001 and *p* < 0.001, respectively with 10,000 permutations). CRN genes were found to have mean 3’ IGDs greater than that observed for non-CRN genes (*p* = 0.0148, with 10,000 permutations), but this was not the case for 5’ regions. The larger IGD in the 3’ but not 5’ region of CRN genes compared to the 5’ region was further investigated by looking at functional annotations of the 5’ neighbouring genes to CRNs. Fifteen of the 34 5’ neighbours of CRN genes were found to have functional annotations, but no clear trend in gene function could be determined. However, not all effector candidates showed these patterns, with no significant difference observed in IGD between protease inhibitors and neighboring genes (*p* > 0.05). Secreted *P. cactorum* CAZymes proteins were found to have significantly greater 5’ IGD. Non-effector candidate elicitins had IGDs with no difference in distribution to all genes (*p* > 0.05). Interestingly, putative non-secreted CAZYmes were observed to have significantly shorter 5’ and 3’ IGDs than the total gene set (*p* = < 0.001 and *p* = < 0.001, respectively with 10,000 permutations). This indicates that the forces driving genomic arrangement of regions containing RxLR and CRN cytoplasmic effector candidates and apploastic CAZyme effector candidates are distinct to those of other effector families in *P. cactorum*.

**Figure 3:**
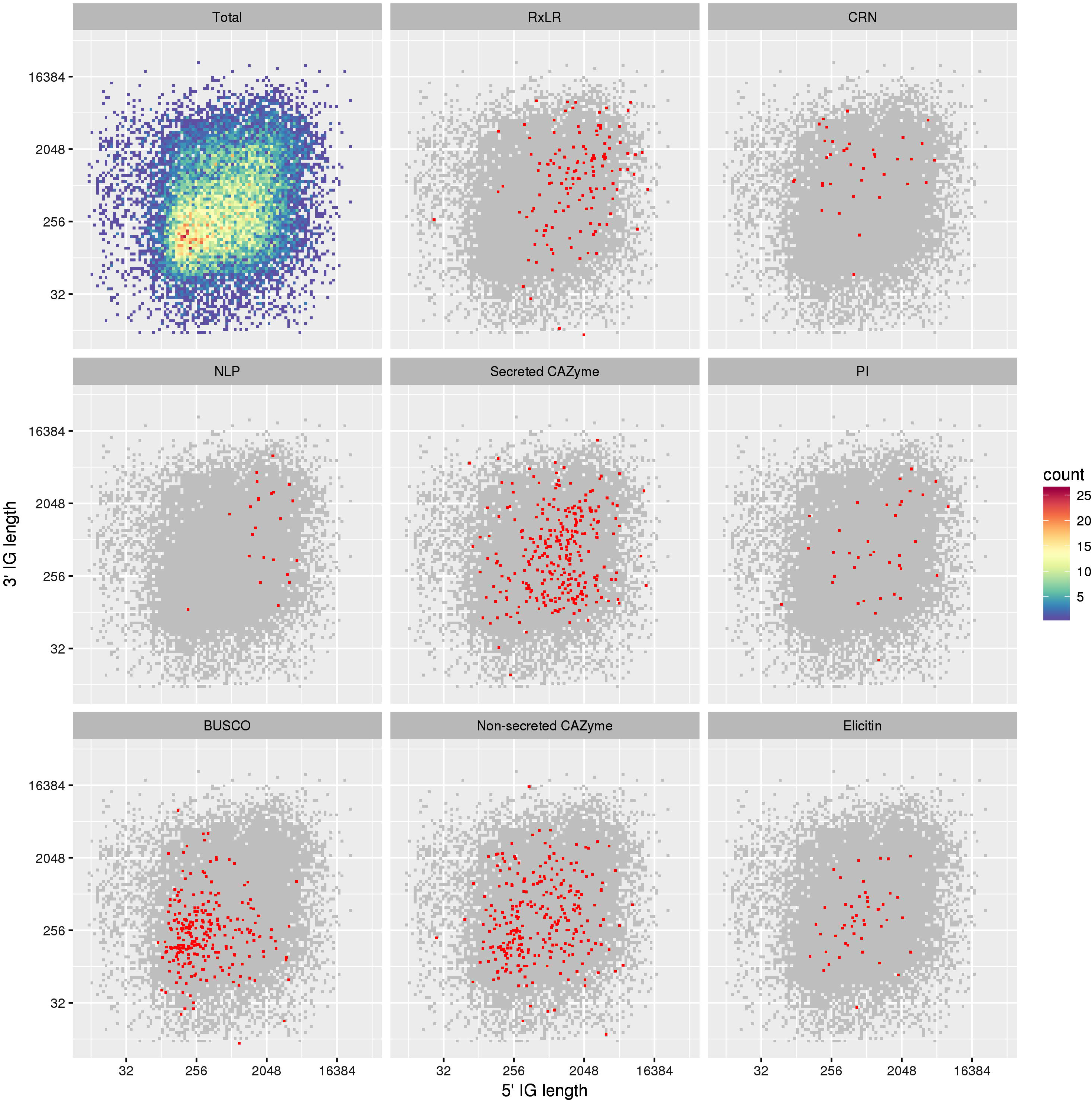
Intergenic distance of cytoplasmic and apoplastic effectors as well as non-effector candidates. Intergenic distance (5’ and 3’) of all *Phytophthora cactorum* 10300 genes is displayed in a density plot (Total) with scale bar indicating gene density within the plot. Additional plots highlight subsets of effector candidates within the distribution including RxLR and Crinkler (CRN) cytoplasmic effector candidates, secreted Carbohydrate-Active enZymes (CAZymes), protease inhibitors and necrosis inducing protein (NLP) apoplastic effector candidates. Distribution of non-effector candidates is shown for conserved eukaryotic genes (BUSCO), non-secreted CAZymes and elicitins.

**Table 6.** Number of genes neighboring the start or end of 4,623 *Phytophthora cactorum* contigs by effector category. The occurrence of genes neighboring contig breaks was not evenly distributed between gene categories (X^2^ = 104.23, df = 8, *p* < 0.01).

|  | Total<br>genes | Neighboring<br>contig breaks | % neighboring<br>contig breaks |
| --- | --- | --- | --- |
| All genes | 23884 | 5041 | 21.1 |
| RxLRs | 199 | 61 | 30.7 |
| CRNs | 76 | 39 | 51.3 |
| NLPs | 24 | 7 | 29.2 |
| Protease inhibitors (all) | 22 | 8 | 36.4 |
| Secreted CAZymes | 282 | 57 | 20.2 |
| Non-secreted CAZY | 410 | 61 | 14.9 |
| Elicitins | 47 | 10 | 21.3 |
| BUSCO genes | 272 | 16 | 5.9 |

## Discussion

### A new genomic resource to study strawberry crown rot

*P. cactorum* is a persistent pathogen of strawberry and an economically significant pathogen of apple (Erwin and Ribeiro 1996). Genomic resources are available for these hosts (Antanaviciute et al. 2012; Hirakawa et al. 2014; Davik et al. 2015; Daccord et al. 2017), and recent work has identified resistance-associated QTL for cultivated strawberry (Nellist et al. 2018). Despite this, genomic resources for the pathogen are limited to identification of ESTs expressed during infection (Chen et al. 2011) and transcript expression during oospore germination characterized (Chen et al. 2014; Chen et al. 2017). We report the sequencing, annotation and assembly of the *P. cactorum* genome, totalling 59 Mb, with a total of 23,884 predicted transcripts. The assembly was fragmented in 4,623 contigs, with 2,913 contigs over 1 kb in length. However, BUSCO statistics were indicative of a highly-complete assembly and detection of 89% of CEGs as present in a single copy within predicted gene models was greater than that observed from other *Phytophthora* spp. Assembly fragmentation was attributed to the high repeat content (18%) observed in the assembly. The level of repetitive content was similar to that observed in the similarly sized genome of *P. sojae* but did not show the same levels of genome expansion as Clade 1 species *P. parasitica* or *P. infestans*. The sequenced and annotated *P. cactorum* genome is an important genomic resource that will aid functional study of effector gene candidates, as well as providing a resource to study the genomic basis of host specificity, which has been reported in the pathogen (Hantula et al. 1997; Lilja et al. 1998; Hantula et al. 2000; Thomidis 2001; Thomidis 2003; Bhat et al. 2006).

### Genomic characterisation of a broad profile of MAMPs and effectors

*Phytophthora* pathogens utilise a diverse range of secreted apoplastic and cytoplasmic effectors to aid infection. This work characterised the *P. cactorum* genome, identifying both apoplastic and cytoplasmic effector candidates as well as non-effectors that are typical of MAMP elicitors of host defence. This study unveiled the diversity of effectors in the *P. cactorum* genome, supplementing those effectors identified during development and cyst germination (Chen et al. 2011; Chen et al. 2014) with those that may be specifically expressed during infection and the transition to necrotrophy. This study identified considerably greater numbers of CRN, elicitins, GH, PL and RxLR candidates than previously identified in the *P. cactorum* transcriptome (Chen et al. 2014). Equal or greater numbers of genes encoding NLPs, protease inhibitors, cutinases and PcF domain-carrying proteins were identified, however some of the candidates were discarded due to possession of a transmembrane domains or a GPI anchor.

This study reports a novel method for CRN prediction. The two-model LFLAK-DWL approach ensures identification is based upon the characteristic N-terminal domains of CRNs and not upon the variable C-terminal functional domains or upon regular-expression searches for conserved motifs, which may not be flexible enough to allow for sequence variation. This provides new opportunities for identification of new functional CRN domains and will advance research in this poorly understood effector family.

### Identification of homologs to well characterised avirulence genes

Establishing orthology between predicted proteomes is an important tool for translation of functional research from model *Phytophthora* species into *P. cactorum*. A total of 20,157 (84%) of predicted *P. cactorum* proteins had identified orthologs in other *Phytophthora* spp. Proteins in shared ortholog groups between *P. infestans* and *P. cactorum* allowed identification of two *Avr1* homologs, one *Avr3b*, one *Avr4*, two *Avrblb1* homologs (of which one was truncated) and two homologs of *AvrSmira2* (Table 5). These characterised avirulence genes represent key targets for further functional study.

### Evidence for a two-speed genome in *P. cactorum*

Effector genes have previously been characterised as showing uneven distributions throughout *Phytophthora* genomes, with measurements of IGD showing that effector genes are located in gene-sparse regions of the *P. infestans genome* (Haas et al. 2009). This has led to the concept of a two-speed genome in these organisms, where different regions of the genome are subject to different evolutionary pressures (Dong et al. 2015). *P. cactorum* RxLR, CRN and secreted CAZyme effector candidates showed increased IGD over non-effector genes, supporting the concept of a two-speed genome in *P. cactorum*. Fragmentation of *P. cactorum* assembly meant that 21% of genes were excluded from this analysis, due to being located on the end of a contig. Unsurprisingly, functional groups of genes within this group were not evenly represented on contig ends with 30% of RxLR and 50% CRN genes located on contig ends in contrast to 6% of BUSCO conserved eukaryotic genes. A high frequency of contig breaks was observed in the 3’ region of CRN genes and may have biased these distances to be shorter than if measurements were taken from a more contiguous assembly. These analysis should be repeated when improved assemblies become available. Furthermore, the low occurrence of conserved eukaryotic genes neighbouring contig breaks highlights that although these genes are comparatively useful in assessing assembly quality, their lack of an even distribution throughout difficult-to-assemble regions means that these genes do not accurately reflect the true “gene-space” in the assembly.

### Outcomes for breeding durable disease resistance

A broad complement of effectors and *Avr* genes are described in our characterisation of the *P. cactorum* genome. Qualitative resistance to *Phytophthora* pathogens is frequently determined by recognition of an RxLR in a gene-for gene dependant manner (Anderson et al. 2015). However, recognition of the *P. infestans* RxLR effector AvrSmira2 in field conditions is associated with quantitative resistance in potato (Rietman et al. 2012). Quantitative resistance to *Phytophthora* diseases has also been associated with basal defence (Vega-Sánchez et al. 2005; Thomas et al. 2007). Accordingly, this study characterises a broad range of effector genes and provides candidates to investigate the basis of quantitative strawberry resistance to *P. cactorum* (Denoyes-Rothan et al. 2004; Shaw et al. 2006; Shaw et al. 2008; Nellist et al. 2018). RxLR effectors are still priority candidates disease related pathogen genes for functional study of strawberry resistance to *P. cactorum*, particularly homologs of *AvrSmira2* characterised avirulence genes.

**Accession number(s)**. The whole-genome shotgun project has been deposited at DDBJ/ENA/GenBank under the accession no. MJFZ00000000. The version described in this paper is version MJFZ01000000.

## ACKNOWLEDGMENTS

ADA, CFN, LAL and RJH were funded by BBSRC grant BB/K017071/1. MBB, EL and sequencing costs were financed by a strategic NIBIO project (basic funding). All authors thank Sophien Kamoun for valuable support throughout the project.

## Supplementary data

**Supp. Data 1**: Functional annotation of *Phytophthora cactorum* predicted proteins, including information on location, sequence, secretion status, identification as an RxLR, crinkler or CAZyme, orthology information (including orthogroup, number of proteins present in the orthogroup by species and orthogroup contained proteins), BLAST homology information (PHIbase, Swissprot and characterized oomycete avr genes) and identified InterProScan annotations.

**Supp. Data 2**: *Phytophthora cactorum* 10300 genes with homology to known *Phytophthora* effector gene candidates. The orthogroup is shown for the query gene, with numbers of genes in each orthogroup shown for *P. cactorum* (Pcac), *P. parasitica* (Ppar), *P. infestans* (Pinf), *P. capsici* (Pcap) and *P. sojae* (Psoj), as well as functional annotation of each gene. Results showing best tBLASTx hits of all *P. cactorum* genes to a custom database with an E-value < 1x10^-30^.

**Supp. Data 3**: Alignment of proteins from the 16 orthogroups representing necrosis inducing proteins (NLP). Conservation of cytosine sites at alignment positions 624 and 661 identifies these proteins as Type1 NLPs.

**Supp. Data 4**: HMM models used for identification of Crinklers (CRNs) in *Phytophthora spp*. proteins. Two HMM models were made, one for identification of the CRN LFLAK domain and one to the CRN DWL domain.

**Supp. Data 5**: Fasta sequences of predicted Crinkler (CRN) proteins from *P. cactorum* (Pcac), *P. parasitica* (Ppar), *P. infestans* (Pinf), *P. capsici* (Pcap) and *P. sojae* (Psoj) genomes.

